# FLEX: A heparin-binding fusion partner engineered from fibroblast growth factor 1 to enhance protein expression, solubility and purity

**DOI:** 10.64898/2026.03.17.712271

**Authors:** Ravina Mistry, John A. Harris, Dominic P. Byrne, Rachael Morris, Yong Li, Chris W. Theron, Stephen B. Kaye, David G. Fernig, Daniel M. Foulkes

## Abstract

Expression of aggregation-prone, unstable, or cytotoxic recombinant proteins remains a major bottleneck in both academic and industrial research. Although solubility-enhancing affinity tags can improve expression, they often compromise purification stringency, increase construct size, or require additional downstream processing. Here we report FLEX, a compact 15.5-kDa dual-function fusion tag engineered from human fibroblast growth factor-1 (FGF1) that integrates intrinsic protein-stabilising properties with high-affinity heparin binding. Structure-guided computational redesign of the FGF1 scaffold reduced exposed hydrophobic residues, removed flexible protease-susceptible regions, and expanded the electropositive surface while preserving the canonical heparin-binding interface. FLEX exhibits markedly improved thermal and chemical stability relative to wild-type FGF1 together with enhanced heparin affinity, enabling high-stringency washing and improved purity in a single affinity step. We demonstrate the broad utility of FLEX by expressing and purifying a panel of challenging proteins in *Escherichia coli*, including cytotoxic *Pseudomonas aeruginosa* virulence factors that are difficult to obtain in active form. Unexpectedly, FLEX also performed robustly in mammalian expression systems, where transiently expressed FLEX-tagged proteins were recovered at higher yield and purity than with gold standard Myc and Strep tags, including difficult targets such as Tribbles 3 (TRIB3). These findings establish FLEX as a versatile affinity-and-stabilisation tag that improves expression and purification across diverse systems, providing a practical new tool for structural, biochemical, and translational studies of otherwise intractable proteins.

## Introduction

Expression of challenging recombinant proteins remains a persistent bottleneck in both basic research and biopharmaceutical development. Purification is often facilitated by generating protein constructs that incorporate fusion tags that either enhance a protein’s stability or enables affinity purification. For example, solubility-enhancing fusion tags such as thioredoxin A (TrxA), small ubiquitin-like modifier (SUMO), HaloTag (Halo), and N-utilization substance protein A (NusA) are intrinsically soluble proteins that can increase the soluble expression of their fusion partners [1]. In contrast, affinity tags are used to capture and purify proteins using a range of selective affinity matrices. PolyHistidine tags, typically comprised of 6-12 His residues at the N- or C-terminus of a protein, are among the most used affinity tags. They enable rapid enrichment of proteins using metal-chelating resins and elution through competition with high concentrations of imidazole but have limited influence on a protein’s solubility. Maltose binding protein (MBP) and glutathione-S-transferase (GST) are also widely used affinity tags that bind to immobilised maltose and glutathione, respectively, whilst also possessing solubility enhancing properties [2]. However, compared to PolyHis, both MBP and GST exhibit slower binding and elution kinetics for their respective resins. Furthermore, their relatively larger sizes (42 kDa for MBP and 26 kDa for GST) can interfere with protein function or structural characterisation, often necessitating tag removal from the final product, thereby increasing labour and processing time [3]. Each fusion tag offers distinct advantages and limitations, with some tags more suitable for specific applications than others. Consequently, pairing a protein of interest (POI) with the right tag (or combinations of tags) through systematic screening, in addition to optimised construct design and selecting an appropriate expression system, are critical factors that often determine the success of a purification strategy.

Our understanding of biochemical processes has been fundamentally transformed by the engineering of microorganisms for production of recombinant proteins. Owing to its rapid growth, scalability, low cost and genetic tractability, *Escherichia coli* remains, by some margin, the most used laboratory-based protein expression systems. However, it has been estimated that just ∼15 % of eukaryotic proteins are soluble when expressed in *E. coli* in their native forms [4, 5]. Many proteins, including bacterial proteases, multidomain assemblies, and aggregation-prone factors, exhibit poor solubility, low yield, or toxicity when expressed in standard *E. coli* strains. Moreover, *E. coli* systems often lack much of the necessary protein maturation architecture found in eukaryotic cells (including post translational modification [PTM] pathways and specific chaperone systems) resulting in expression of ‘misfolded’ proteins in inactive or unstable forms. In addition to incorporation of a fusion tag, interventions to improve *E. coli* systems have included strain engineering, co-expression of molecular chaperones or enzymes to add PTMs, and low temperature induction conditions [6, 7]. In recent years, alternative expression systems, such as mammalian, insect cell cultures or cell-free systems have also been developed to overcome the contextual limitations inherent to heterologous protein expression in microbes [8, 9]. These systems, which still often employ fusion tags, can more closely replicate the native cellular environment of a eukaryotic protein, and have even enabled production of challenging targets, such as membrane proteins and multi-protein complexes that are typically refractory to standard microbial expression pipelines. However, these advantages are often offset by substantial compromises, most notably reduced yields and concomitant increases in production time and costs. Given these challenges, supplementing the biochemist’s toolbox with novel tools to support the expression and purification of a broader range of protein targets is of considerable importance.

Fibroblast growth factors (FGFs) are archetypal heparin/heparan sulfate binding proteins [10] through affinity directed electrostatic interactions [11, 12]. This feature of FGFs has been utilised for purifying FGF-family proteins using immobilised heparin [13]. Immobilised heparin has therefore been widely used as a high-affinity matrix for purifying FGFs and other HS-binding and nucleic acid binding proteins. Native FGF1 is the ‘universal FGF from the perspective of its broad selectivity for sulfation patterns in heparin [14] and FGF receptors. However, it is relatively unstable and prone to aggregation limiting its utility as a fusion handle [15]. More broadly, the application of naturally occurring heparin-binding proteins as affinity tags has been constrained by their size, folding requirements, and sensitivity to chemical and thermal stress.

Here, we present FLEX, a compact (15.5 kDa) engineered fusion tag derived from human FGF1, designed to simultaneously enhance expression, solubility, stability, and heparin-binding affinity. Using structure-guided engineering and computational design, we introduced targeted amino acid substitutions that reduce exposed hydrophobicity, remove flexible and protease-susceptible regions, and expand the electropositive surface while preserving the core heparin-binding interface. FLEX retains the characteristic β-trefoil fold of FGF1 but exhibits markedly improved thermal and chemical stability.

We demonstrate that FLEX functions as a versatile purification handle for a broad range of challenging proteins, including cytotoxic *Pseudomonas aeruginosa* virulence factors (ExoU, ExoS and LasB). FLEX enables high-yield, single-step purification via heparin affinity chromatography under stringent wash conditions and, in several cases, improves yield and purity compared to conventional workflows, such as immobilised metal affinity chromatography (IMAC).

Unexpectedly, FLEX also proved highly effective in mammalian expression systems, where affinity purification is often limited by low expression levels, complex lysates, and co-purifying host proteins. In transiently transfected HEK293T cells, FLEX increased recovery of soluble protein and enabled efficient purification of difficult targets such as TRIB3, a pseudokinase that has been challenging to obtain in quantities sufficient for biochemical analysis despite extensive efforts using conventional tags. These findings were surprising given that heparin affinity chromatography is rarely used as a general-purpose strategy in mammalian expression workflows, yet they highlight the ability of FLEX to function robustly across diverse cellular contexts.

Collectively, these results establish FLEX as a dual-function affinity and stabilisation tag that overcomes key limitations of conventional fusion handles. By integrating structural stabilisation with high-affinity heparin binding, FLEX provides a modular platform for the expression, purification, and functional characterisation of otherwise recalcitrant protein targets, from bacterial toxins to mammalian signalling proteins, and represents a valuable addition to protein production pipelines.

## Materials and methods

### Sequence analysis and structural modelling

Amino acid sequences (residues 17-152) of WT FGF1 (UniProt: P05230) and the engineered FLEX variant were aligned using standard pairwise alignment tools using the following sequences:

FGF1 WT

NLPPGNYKKPKLLYCSNGGHFLRILPDGTVDGTRDRSDQHIQLQLSAESVGEVYIKSTETGQYLAMDT DGLLYGSQTPNEECLFLERLEENHYNTYISKKHAEKNWFVGLKKNGSCKRGPRTHYGQKAILFLPLPV

FLEX

NLPPGNYNKPQLLYCQNGGYFLRILPDGTVDGTRDENDPYIILQLSAESVGVVYIKSTETGLYLAMDKDGRLYGSKTPNDECLFLERLEENHYNTYISKKYAEKNWFVALKKNGSCKRGPRTHYGQKAILFLPLPV

Sequence visualisation was performed with Jalview 2.11.5.1 (Tcoffee with defaults pairwise alignment) [16] and sequence analysis was conducted with Expasy Tools [17] for hydrophobicity [18]. Amino acid sequences (residues 17-152) of WT FGF1 (UniProt: P05230) and the engineered FLEX variant were aligned (Figure 1A). Structural visualisation of WT FGF1 was based on the published crystal structure (PDB ID: 1JQZ). For surface comparison, a structural model of FLEX was generated with MODELLER version 10.4 (Webb and Sali, 2016). Surface hydrophobicity was coloured in Pymol according to the Eisenberg hydrophobicity scale [19]. Surface electrostatic surface was predicted between +/-range of 3, using the APBS electrostatics plugin in Pymol.

**Figure 1:**
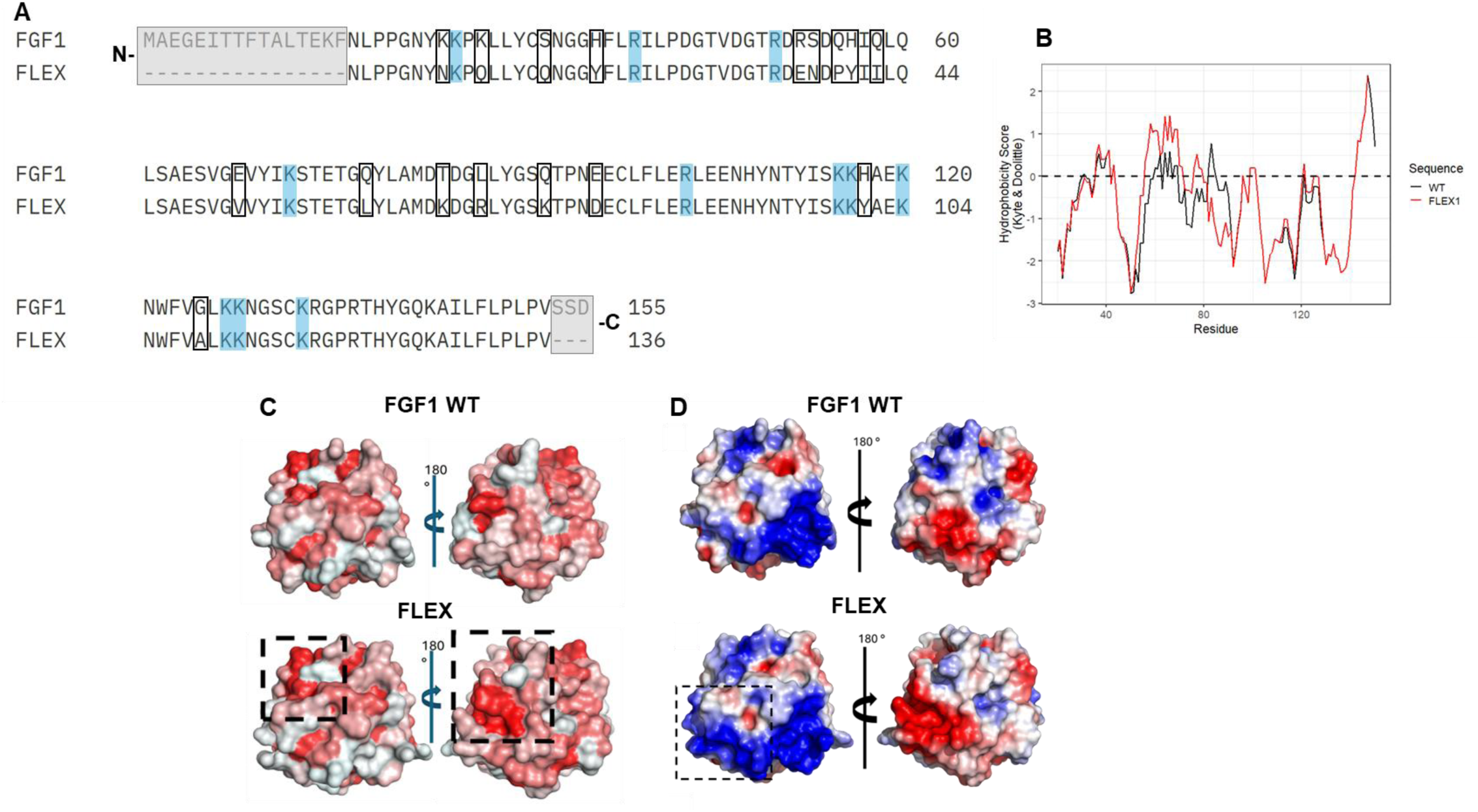
Structural and sequence analysis of FGF1 and the engineered FLEX tag. (A) Sequence alignment of WT FGF1 and engineered FLEX. Lysine (K) and arginine (R) residues contributing to the β-trefoil electropositive surface are highlighted in blue. Engineered stabilising substitutions outside the heparin-binding interface are indicated with black boxes. Truncated residues in FLEX are shown in gray boxes. Numbering corresponds to WT FGF1. (B) Sequence hydrophobicity profiles of WT FGF1 and FLEX, calculated using the Kyte-Doolittle scale (Kyte and Doolittle, 1982). Residues with values > –0.5 are considered hydrophobic (indicative of buried/internal regions), while residues < –0.5 are hydrophilic (indicative of solvent-exposed regions). (C) Surface hydrophobicity predictions of WT FGF1 (PDB ID: 1JQZ) and FLEX (AlphaFold model) generated using Modeller 10.4. Front (heparin-binding) and posterior faces (180° rotation) are shown, where red denotes hydrophobic regions. Enhanced hydrophobic patches in FLEX are indicated with dashed boxes. (D) Surface mapping of heparin-binding residues on WT FGF1 and FLEX. Key Lys/Arg residues are shown in blue. Electrostatic surfaces are displayed, with red indicating negative potential and blue indicating positive potential. The dashed box highlights the expanded positively charged surface in FLEX.

Sequence hydrophobicity profiles were calculated using the Kyte-Doolittle hydropathy scale, where values greater than −0.5 were considered hydrophobic (indicative of buried or internal regions) and values less than −0.5 hydrophilic (indicative of solvent-exposed regions). Structural visualisation of WT FGF1 was based on the published crystal structure (PDB ID: 1JQZ). A structural model of FLEX was generated using AlphaFold2 [20]. Surface hydrophobicity predictions were generated using MODELLER version 10.4, and electrostatic surface potentials were visualised using standard molecular graphics software. Front (heparin-binding) and posterior (180° rotated) surface views were generated for comparative analysis. The FLEX fusion tag construct, encoding an N-terminal 6×His tag followed by FLEX, a TEV protease cleavage site, and a multiple cloning site in the pET-28a(+) vector.

### Plasmids

All bacterial expression constructs were cloned into the pET-28a(+) vector, which encodes an N-terminal hexahistidine tag. For FLEX-containing constructs, the FLEX sequence was inserted downstream of the His tag, followed by a tobacco etch virus (TEV) protease cleavage site and the protein of interest. Mammalian expression constructs were cloned into the pTWIST CMV vector. All DNA sequences were synthesised and sequence-verified by Twist Bioscience prior to use.

### Transformation in *E.* coli

Chemically competent *E.* coli were thawed on ice (10 min), and 80 ng of respective plasmid was added to 50 µL cells. The mixture was placed on ice (10 min), incubated at 42 °C (30 s) in a water bath, and then placed back on ice (2 min). Room temperature Luria broth (LB, Melford) (200 µL) was added to the transformation mixture and incubated at 37 °C for 1 h (200 rpm). Following incubation, 50 µL of culture was spread on to the appropriate selection plate and incubated overnight (37 °C).

### Protein expression and purification in *E. coli*

Recombinant proteins were expressed in *E. coli* BL21(DE3) pLysS cells. Cultures were grown in LB medium supplemented with appropriate antibiotics at 37 °C to mid-log phase before induction with 0.4 mM isopropyl-β-d-thiogalactopyranoside (IPTG) at 18°C for 18 h before harvesting by centrifugation. Cells were resuspended in the indicated lysis buffers (Tris pH 7.4 or 8.2). Lysis buffers also contained 300 mM NaCl, 0.1% (v/v) Triton-X-100, 10 mM imidazole, 1 mM DTT, 10% (v/v) glycerol and a cOmplete protease inhibitor cocktail tablet (Roche). Cells were lysed by sonication using Digital Sonifier SFX 550 at 35% amplitude, 6-10 rounds of 15 s on and 1 min off on ice. Lysates were clarified by centrifugation (15,000 x g for 60 min) to remove insoluble material prior to chromatographic purification.

### Immobilised metal affinity chromatography (IMAC)

His-tagged proteins were purified using nickel-nitrilotriacetic acid (Ni-NTA) resin packed into 1 mL columns (Cytiva/Bio-Rad). Clarified lysates were applied to equilibrated columns, washed with 20 column volumes of wash buffer (20 mM Tris, 300 mM NaCl and 30 mM imidazole) to remove non-specifically bound proteins, and eluted using buffer containing 500 mM imidazole with 10% (v/v) glycerol. Elution fractions were collected in 500 µL volumes. Where indicated, IMAC eluates were subjected to further purification by size-exclusion chromatography. Protein concentrations were determined using the Bradford assay (ThermoFisher) according to the manufacturer’s instructions, with bovine serum albumin (BSA) used as a standard. Concentrations were independently estimated by measuring UV absorbance at 280 nm using a NanoDrop spectrophotometer.

### Heparin affinity chromatography (HAC)

Heparin affinity chromatography was performed using 1 mL immobilised heparin columns (Cytiva) equilibrated in 20 mM Tris buffer with 300 mM NaCl. Clarified lysates were loaded onto the column and washed with buffer containing either 300 mM or 1 M NaCl to remove non-specifically bound proteins. Bound proteins were eluted either stepwise or using linear NaCl gradients up to 2 M NaCl, with 500 µL fractions collected. Protein elution was monitored by absorbance at 280 nm or Bradford assay.

### Size-exclusion chromatography (SEC)

Size-exclusion chromatography was performed using a Superdex 200 16/600 column (Cytiva) equilibrated with two column volumes of 20 mM Tris-HCl (pH 7.4) containing 100 mM NaCl. Chromatography was conducted at a flow rate of 0.5 mL min⁻¹ using an ÄKTA chromatography system. Purified proteins were analysed to assess oligomeric state and aggregation prior to downstream biophysical assays. Elution was monitored by absorbance at 280 nm, and fractions were collected automatically.

### Isothermal titration calorimetry (ITC)

FGF1 and FLEX were purified to homogeneity by heparin affinity chromatography followed by SEC. Freeze dried polymeric heparin (14,000 Da) was resuspended in SEC buffer (20 mM Tris pH 7.4, 100 mM NaCl, 10% v/v glycerol), were measured using a MicroCal PEAQ-ITC instrument. Experiments were conducted at 25 °C in Tris buffer (pH 7.4) containing 100 mM NaCl. Polymeric heparin was titrated into protein solutions using a total of step-wise 19 injections, and the heat energy change accompanying the reaction was detected by comparison with a reference cell. The Heat changes were integrated and fitted using a single-site binding model supplied with the MicroCal PEAQ-ITC software. Representative thermograms from three independent experiments are shown, and dissociation constants (K_D_) are reported as mean ± standard deviation.

### Differential scanning fluorimetry (DSF)

Thermal stability of WT FGF1 and FLEX was assessed by DSF using an Applied Biosystems StepOnePlus Real-Time PCR instrument. Proteins were diluted to 5 µM in 20 mM Tris-HCl (pH 7.4), 100 mM NaCl, and 1 mM DTT, and incubated with the indicated concentrations of urea in a total reaction volume of 25 µL. SYPRO Orange (Invitrogen) was used as the fluorescent probe. Fluorescence at 570 nm was monitored while the temperature was increased from 25 °C to 95 °C in 0.5 °C increments. Melting temperatures (Tm) were determined from the inflection point of the unfolding transition and average *T*_m_/Δ*T*_m_ values were calculated as previously described [21] using the GraphPad Prism software. Each experiment was performed in triplicate, with each condition assayed in technical duplicate.

### Enzymatic activity assays

Enzymatic activities of purified *P. aeruginosa* virulence factors were measured in 96-well plate format using a Hidex Sense plate reader. All assays were performed at a final volume of 50 µL with enzymes at 100 nM concentration. Where indicated, the FLEX tag was removed by TEV protease cleavage followed by reverse affinity purification prior to analysis. ExoU phospholipase activity was measured using arachidonyl thio-phosphatidylcholine as substrate in the presence of 1 µM phosphatidylinositol-4,5-bisphosphate (PIP₂, Avanti polar lipids) and 5 µM mono-ubiquitin. Hydrolysis was monitored using 5,5′-dithiobis-(2-nitrobenzoic acid) (DTNB) as a reporter by measuring the increase in absorbance at 405 nm [22, 23]. ExoS ADP-ribosyltransferase activity was assessed by ExoS auto-ADP-ribosylation using ε-NAD (25 µM, Merck) as the substrate and 5 µM 14-3-3β as the ExoS activating co-factor.

Fluorescence was monitored using an excitation wavelength of 300 nm and emission at 410 nm [23, 24]. LasB and ArpA protease activity was measured using the fluorogenic peptide substrate Mca-KPLGL-Dpa-AR-NH₂ (25 µM, Biotechne) in Tris buffer (pH 8.2) containing 100 mM NaCl, 10 µM ZnSO₄, and 100 µM CaCl₂. Fluorescence was monitored using excitation at 355 nm and emission at 405 nm. All assays were performed with two technical replicates for each condition and three independent experiments.

### Mammalian expression and purification

The kinase domain of protein kinase A (PKA) was cloned into a pTWIST CMV mammalian expression vector with an N-terminal His-FLEX tag. HEK293T cells, cultured in Dulbecco’s modified Eagle medium (Lonza) supplemented with 10% fetal bovine serum (HyClone), penicillin (50 U/ml), and streptomycin (0.25 µg/ml) (Lonza) and maintained at 37°C in 5% CO_2_ humidified atmosphere, were seeded in 10 cm dishes and at 40-50% confluence and transfected using a 3 : 1 polyethylenimine (PEI [branched average *M*_w_ ∼25 000 Da; Sigma-Aldrich]) to DNA ratio (30 : 10 µg, for a single 10 cm culture dish). Proteins were harvested 48 h post transfection in a lysis buffer containing 50 mM Tris–HCl (pH 7.4), 150 mM NaCl, 0.1% (v/v) Triton X-100, 1 mM DTT, 0.1 mM ethylenediaminetetraacetic acid (EDTA), 0.1 mM ethylene glycol-bis(β-aminoethyl ether)-*N*,*N*,*N*′,*N*′-tetraacetic acid (EGTA) and 5% (v/v) glycerol and supplemented with a protease inhibitor cocktail tablet and a phosphatase inhibitor tablet (Roche). Lysates were briefly sonicated on ice and clarified by centrifuged at 20 817×***g*** for 20 min at 4°C, and the resulting supernatants were split in half and equal volumes incubated with 20 µl of either Ni-NTA or immobilised heparin resin (as described above) for 3 h with gentle agitation at 4°C. Affinity beads containing bound protein were collected and washed four times in 1 ml 50 mM Tris-HCl (pH 7.4) and 500 mM NaCl and purified proteins were then eluted from the suspended beads with 30 µl elution buffer containing 50 mM Tris–HCl (pH 7.4) and 500 mM imidazole (with 100 mM NaCl) or 1 M, 1.5 M or 2 M NaCl. Eluted fractions were analysed SDS-PAGE and western blotting and protein detected Ponceau staining and Western blotting using an anti-6×His antibody (Cell Signalling Technologies).

### SDS-PAGE and Western blotting

Samples were heated for 5 min at 95 °C in sample buffer (50 mM Tris–Cl pH 6.8, 1% (w/v) SDS, 10% (v/v) glycerol, 0.01% (w/v) bromophenol blue, and 10 mM DTT). Samples were resolved by SDS-PAGE and either visualised by Coomassie staining or transferred to nitrocellulose membranes (Bio-Rad) by western blotting. Membranes were blocked in Tris-buffered saline with 0.1% (v/v) Tween 20 (TBS-T) in 5% (w/v) non-fat dried milk (pH 7.4) followed by incubation with primary anti-6His tag antibody (Bio-Rad or Cell Signalling Technologies) and secondary horse radish peroxidase-conjugated anti-mouse (ThermoFisher) antibody. Proteins were detected using chemiluminesence reagent (Bio-Rad) and visulised using a Bio-Rad ChemiDoc.

## Results

### Rationale and design of the FLEX mutations

Wild-type (WT) human FGF1 was chosen as a scaffold due to its well-characterised β-trefoil fold and its interaction with heparin and heparan sulfate, which displays relatively low selectivity for sulfation patterns, binding oligosaccharides containing any pair of N-, 6-*O*-, or 2-*O*-sulfate groups [25]. Moreover, the FGF1-heparin binding interface has been extensively characterised by biochemical, mutational, and structural studies [12]. Key features of FGF1 on analysis of high-resolution crystal structure (PDB: 1JQZ [26]) include a positively charged acidic patch mediating glycosaminoglycan binding as well as adjacent regions of exposed hydrophobicity. Leveraging these structural insights in combination with predictions generated by Protein Repair One Stop Shop (PROSS, [27]), we introduced targeted modifications aiming to improve stability and retain heparin affinity of FGF1. Sequence alignment of FGF1 and the engineered FLEX highlights the preservation of basic residues contributing to heparin binding and electropositive surface (blue, Figure 1A) and subsequent substitutions for stabilisation (black boxes, Figure 1A). In addition, FGF1 was truncated to remove N-terminal residues 1-16, which are disordered and dispensable for heparin binding, and the three C-terminal residues (SSD), which are distal to the canonical heparin-binding site (grey, Figure 1A). Importantly, all Lys and Arg residues directly mediating heparin contact were preserved, and no substitutions were introduced within the core motifs.

#### Analysis of Surface properties for FGF1 and FLEX

Surface and Kyte-Doolittle hydropathy analysis (Figure 1C & 1B) of the FLEX and FGF1 sequences revealed several regions of altered hydrophobicity and charge in the engineered protein. Hydrophobicity of each residue was calculated and visualised onto the structural models (Figure 1C). WT FGF1 was predominantly a hydrophilic (Figure 1C, top panels), neutral charged heparin-binding face (Figure 1D, top left panel). Electrostatic surface mapping of FLEX (Figure 1D, bottom panels) displayed extended positively charged patch adjacent to the conserved heparin-binding site (Figure 1D, bottom left, black box). By contrast, the opposing posterior surface provides a more negative (Figure 1D, bottom right panel) and contiguous hydrophobic patch (Figure 1C, bottom left panel). This asymmetric distribution of surface properties may contribute to the stabilising effect of FLEX when fused to aggregation-prone partners explored further in the expression studies described above.

Altogether, these biochemical and structural features from the modification in sequence, suggest that the improved stability and functional properties of FLEX arise through improved core folding, reduced exposed hydrophobic surface area (Figure 1C), and optimised electrostatic environment around the heparin-binding interface (Figure 1D); without altering residues mediating ligand contact. These features provide a foundation for its use as a high-stringency heparin-binding fusion tag.

### Biochemical characterisation of FLEX

#### Expression and purification of WT FGF1 and FLEX

To assess the performance and combinatorial effect of multi-site variation, the yields from bacterial expression between WT FGF1 and the engineered FLEX tag were compared, utilising constructs with an N-terminal hexa-histidine tag expressed in *E. coli*. Purification of His-FGF1 with both immobilised nickel affinity chromatography (IMAC) (Figure 2A, left) and heparin affinity chromatography (HAC) (Figure 2B, left) resulted in lower recovery, consistent with poor tractability and yield of WT FGF1 in bacterial expression systems.

**Figure 2:**
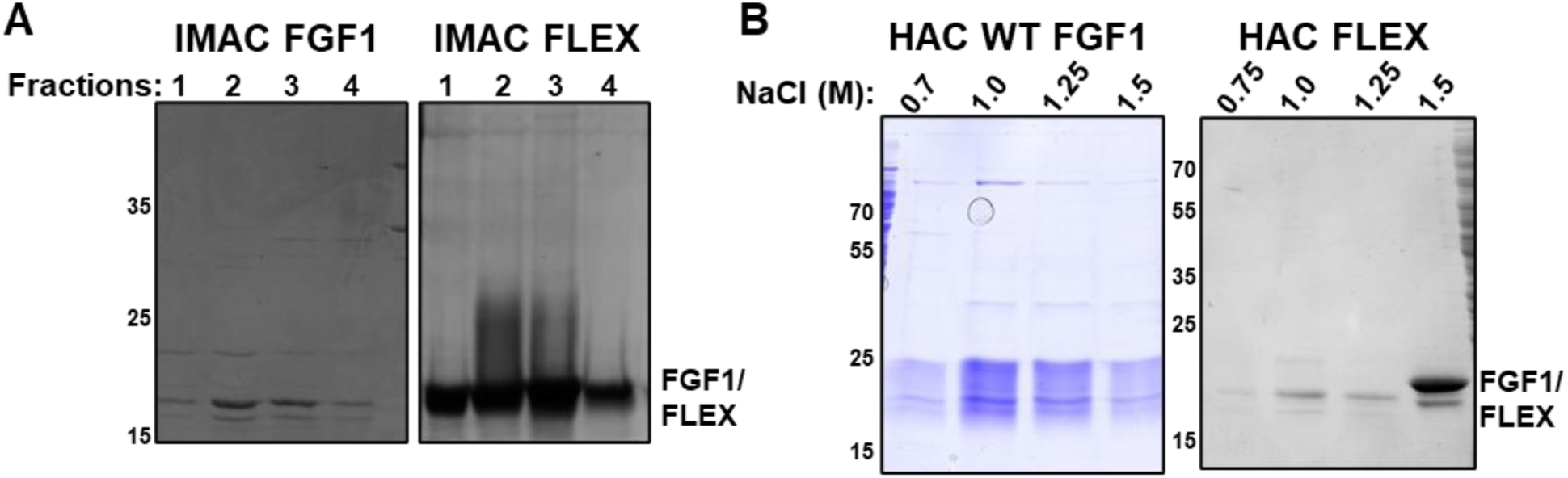
Expression and purification of His-tagged WT FGF1 and FLEX from E. coli. (A) Immobilised metal affinity chromatography (IMAC) purification of His-tagged WT FGF1 (left) and His-FLEX (right) expressed in 2 L *E. coli* cultures. Lanes 1-4 represent elution fractions collected in 500 µL volumes using 500 mM imidazole. (B) Heparin affinity chromatography (HAC) purification of His-tagged WT FGF1 (left) and His-FLEX (right). Following washing with 700 mM NaCl, proteins were eluted in 500 µL fractions at the indicated NaCl concentrations.

While higher purity was observed with the isolation of His-FLEX-tag at 1.5 M NaCl with HAC (Figure 2B, right, Lane 5), in part due to higher heparin binding, there was also a substantial improvement in recovery of the His-FLEX-tag when used in conjunction with IMAC (Figure 2A, right). Accordingly, yields increased from ∼0.5 mg L⁻¹ of *E. coli* culture for FGF1 to ∼4 mg L⁻¹ for FLEX following IMAC purification. These data indicated that the engineered mutations improved expression and yield; and were suggestive of improved heparin affinity, relative to WT FGF1. To assess oligomeric state and aggregation propensity, purified proteins were analysed by size-exclusion chromatography (SEC). Both His-WT FGF1 and His-FLEX eluted as single symmetrical peaks corresponding to monomeric species (Supplementary Figure 1).

#### FLEX exhibits higher affinity for heparin than FGF1

Heparin affinity chromatography was used to establish the chromatographic properties of FGF1 and FLEX, which were compared using a linear NaCl elution gradient. Clarified lysates were loaded onto heparin columns, washed with 700 mM NaCl to remove non-specific interactions, and eluted with a 300 mM-3 M NaCl gradient. FLEX exhibited greater heparin binding as it eluted at higher salt concentrations (∼1.75 M NaCl), In contrast, WT FGF1 eluted at 1.25 M NaCl (Supplementary Figure 2)., which is consistent with its established heparin affinity purification [13]. These data were then used to establish a simple stepwise column washing and elution for FLEX. Washing the column in 1 M NaCl afforded complete removal of contaminants, and elution at 2 M NaCl would recover the FLEX. It is interesting to note that there is a lower molecular weight band apparent in the elution. This is the consequence of the ^31^DG^32^ motif, which is often partially cleaved upon boiling of a protein in Laemmli sample buffer [28, 29].

To quantitatively assess the interaction of FLEX with heparin, we performed isothermal titration calorimetry (ITC) using heparin (∼14,000 Da). FGF1 bound heparin with an apparent dissociation constant (K_D_) of 3.6 µM (Figure 3A), which is consistent with the values obtained for the interaction with a variety of heparan sulfates in an optical biosensor 0.4 µM-8.9 µM [30] and by ITC (1.3 µM, [31]) By contrast, FLEX displayed an enhanced affinity with a K_D_ of approximately 0.5 µM (Figure 3B). The relatively low affinities reflect the fact that although a 1:1 model fitted the data, heparin is polydisperse and polyvalent, so the K_D_ will be an average of high and low affinity binding sites, the latter including ones that may be partly sterically hindered by an already bound FLEX. These data confirm that the engineered mutations strengthen heparin binding without disrupting the canonical recognition site. Based on our structural analysis (Figure 1), the improved binding properties of FLEX arise from a broader electropositive surface rather than changes to the primary binding residues, providing a foundation for its use as a high-stringency heparin-binding fusion tag.

**Figure 3:**
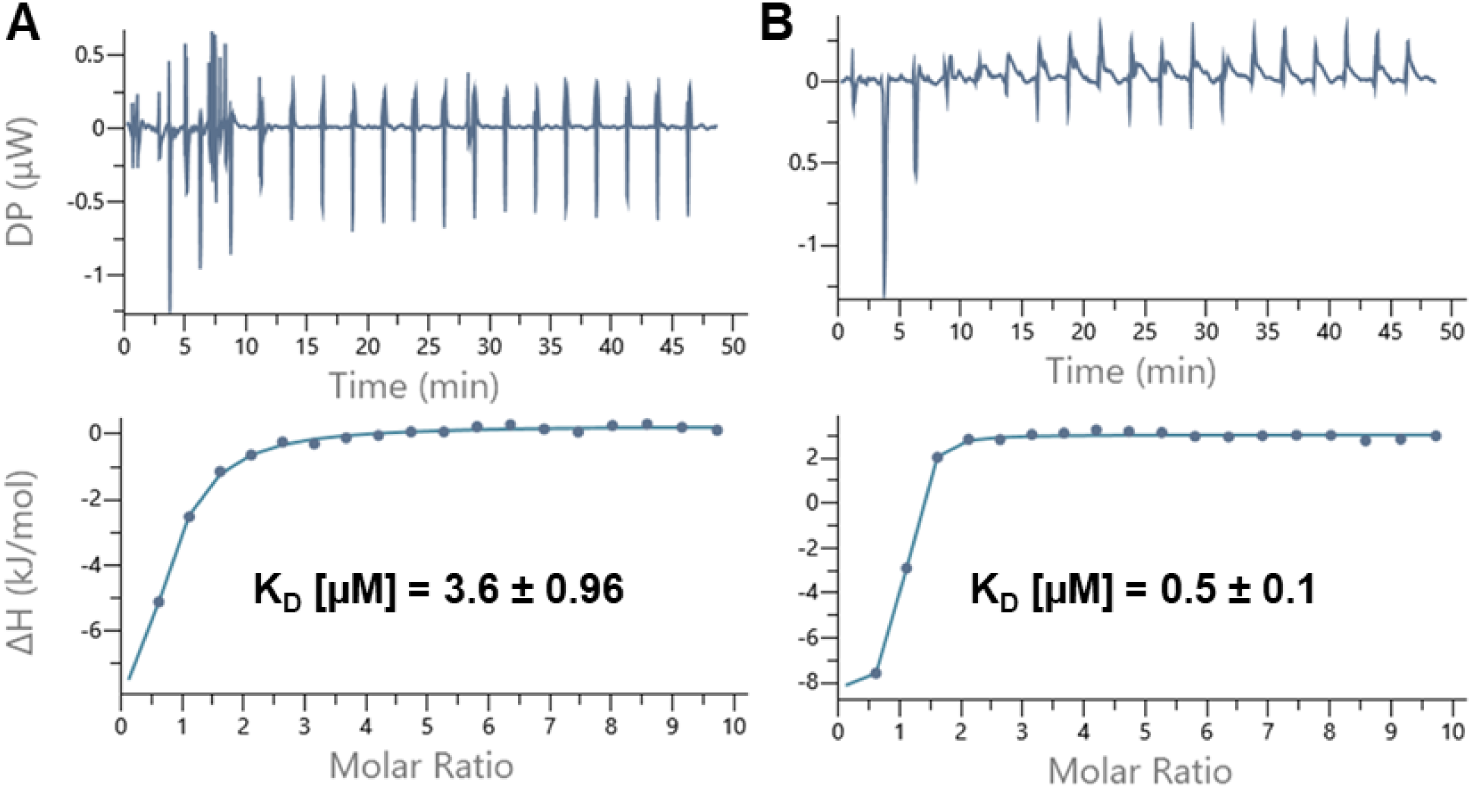
Isothermal titration calorimetry (ITC) analysis of heparin binding to FGF1 and FLEX. Representative ITC thermograms (top panel) and corresponding integrated binding isotherms (bottom panel) for (A) WT FGF1 and (B) FLEX titrated with polymeric heparin from three independent experiments. Experiments were performed at 25 °C in Tris pH 7.4 with 100 mM NaCl. Data were fitted using a single-site binding model.

#### Thermal and chemical stability of FLEX

We next assessed the thermal stability of FGF1 and FLEX using differential scanning fluorimetry (DSF). FGF1 exhibited a melting temperature (T_m_) of 52.1 ± 1.4 °C, whereas FLEX displayed a substantially higher T_m_ of 74.1 ± 1.1 °C (Figure 4A), representing a 22.0 °C increase in thermal stability.

**Figure 4:**
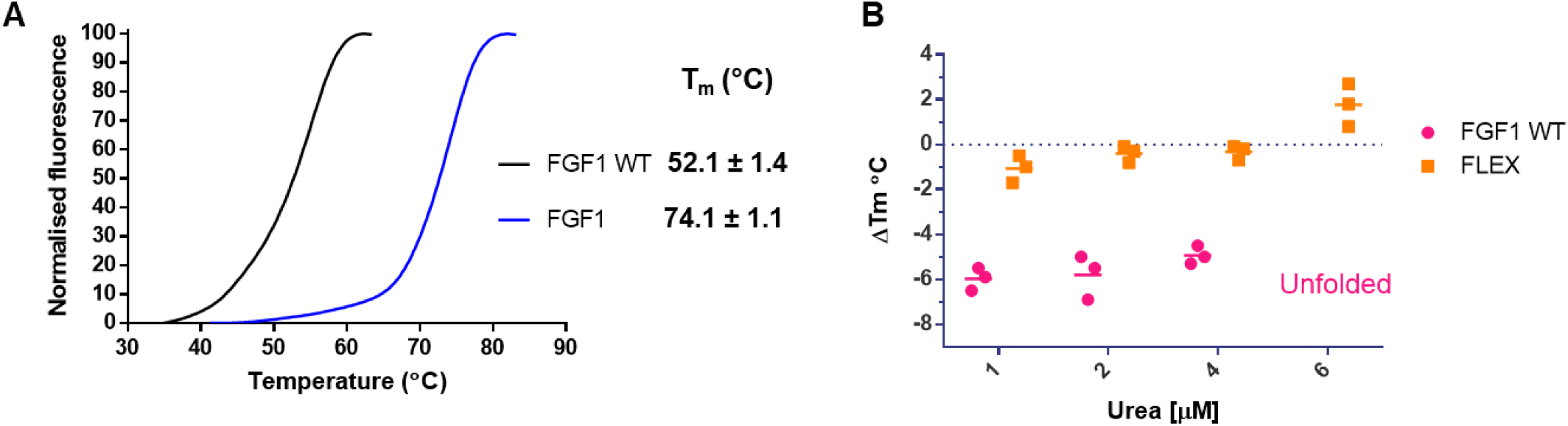
Differential scanning fluorimetry (DSF) analysis of WT FGF1 and FLEX. (A) Thermal denaturation profiles of recombinant WT FGF1 and FLEX. A representative unfolding trace is shown. Melting temperatures (T_m_) are reported as mean ± SD. (B) Change in melting temperature (ΔT_m_) for WT FGF1 and FLEX in the presence of increasing concentrations of urea (1-6 M), expressed relative to the corresponding protein in the absence of urea. All DSF measurements were performed using 5 µM protein from three independent experiments, with each condition assayed in duplicate.

To further evaluate robustness under denaturing conditions, DSF measurements were performed in the presence of increasing concentrations of urea (1-6 M) (Figure 4B). FGF1 WT was highly sensitive to chemical denaturation, exhibiting a change in T_m_ (Δ T_m_) of −6.0 ± 1.8 °C in the presence of just 1 M urea. In 6 M urea, FGF1 WT was completely unfolded (Supplementary Figure 3A). In striking contrast, FLEX retained thermal stability even at 6 M urea, with no significant reduction in melting temperature observed (Figure 4B and Supplementary Figure 3B). These results demonstrate that FLEX is both thermally and chemically stabilised relative to WT FGF1.

### Production of difficult-to-express proteins using FLEX

To assess the utility of FLEX as a fusion partner for challenging recombinant proteins, purification yield and recovery was compared to existing affinity chromatography tools. We first examined expression of ExoU, a *P. aeruginosa* phospholipase and validated virulence factor [32]. His-tagged ExoU purified by IMAC was co-purified with multiple contaminating proteins of similar molecular weight (Figure 5A, panel 1), which were previously identified by mass spectrometry as bifunctional polymyxin resistance protein ArnA and glutamine-fructose-6-phosphate aminotransferase [22]. These have previously been documented as contaminants in IMAC purifications from *E. coli* [33]. Fusion to maltose-binding protein (MBP) or glutathione S-transferase (GST) failed to yield detectable ExoU, with purification primarily recovering the cleaved fusion tags rather than full-length protein (Figure 5A). In contrast, expression of FLEX-ExoU enabled recovery of full-length protein, to relatively high purity, following a single heparin affinity chromatography step (Figure 5A, panel 4).

**Figure 5:**
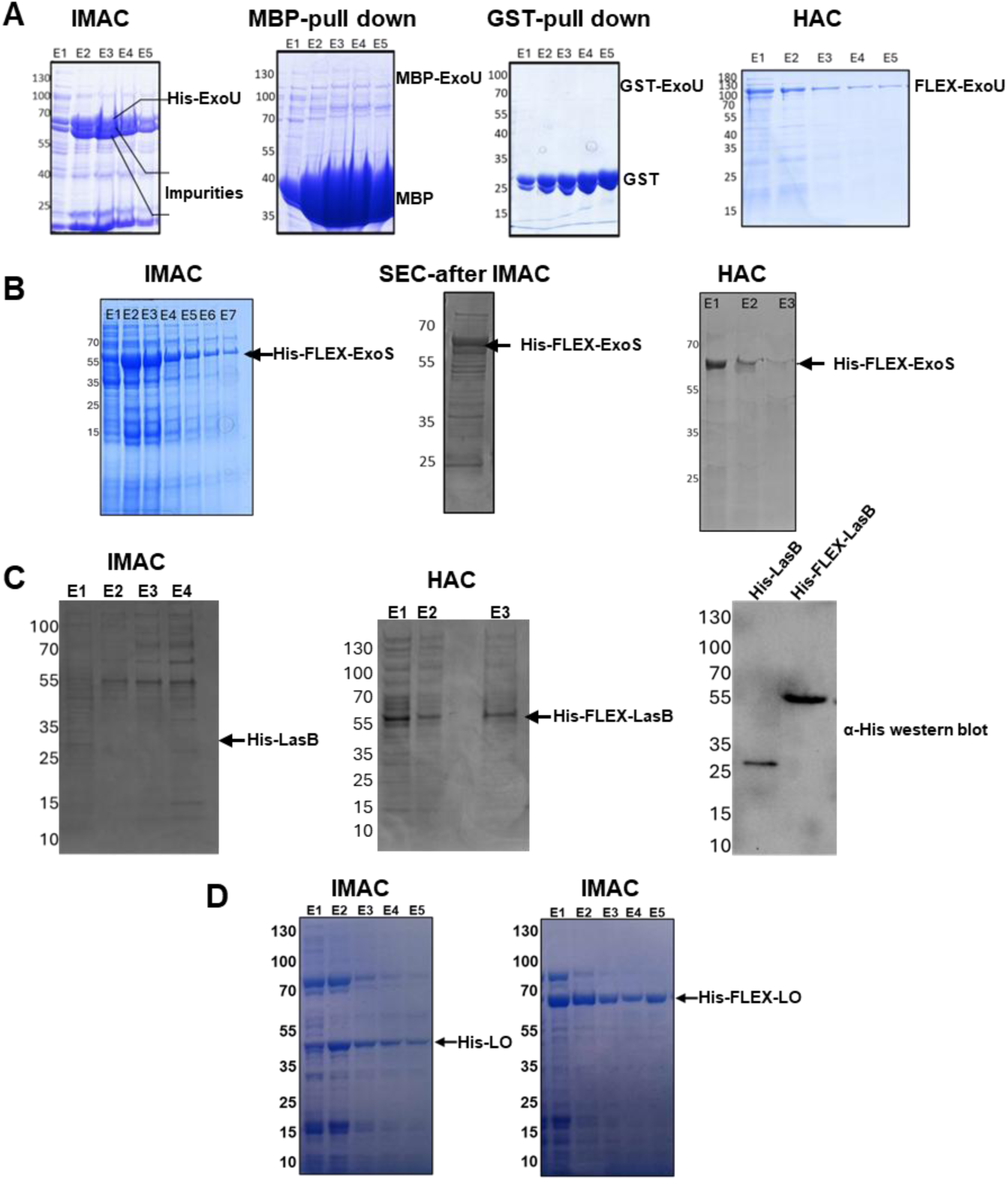
Production of challenging enzymes using FLEX. (A) SDS-PAGE analysis of ExoU expressed with different fusion tags. ExoU was expressed in BL21(DE3)pLysS *E. coli* (2L) as an N-terminally His-tagged construct or fused to maltose-binding protein (MBP), glutathione S-transferase (GST), or FLEX. His-tagged ExoU was purified by immobilised metal affinity chromatography (IMAC), with elution performed using 500 mM imidazole and 500 µL fractions collected. MBP- and GST-tagged ExoU were purified using the corresponding affinity resins, with elution by maltose or reduced glutathione, respectively. His-FLEX-ExoU was purified by heparin affinity chromatography (HAC) using a 1 mL heparin column, washed with buffer containing 1 M NaCl, and eluted with 2 M NaCl in 500 µL fractions. All purification steps were performed in Tris buffer (pH 8.2). (B) SDS-PAGE analysis of His-FLEX-ExoS purified from Escherichia coli. His-FLEX-ExoS was expressed in 2 L cultures and purified using either IMAC or HAC, each performed using a 1 mL chromatography column. IMAC elution fractions (500 µL) were collected using 500 mM imidazole (left), and the eluate was subsequently analysed by size-exclusion chromatography (SEC) (right). For HAC, clarified lysate was washed with buffer containing 1 M NaCl prior to elution with 2 M NaCl, with 500 µL fractions collected. (C) SDS-PAGE and Western blot analysis of LasB expressed with His or His-FLEX fusion tags. His-LasB was expressed in Escherichia coli and purified by IMAC using a 1 mL nickel column, with elution performed using 500 mM imidazole and 500 µL fractions collected (left). His-FLEX-LasB was expressed and purified by HAC (middle). Western blot analysis of IMAC-purified His-LasB and HAC-purified His-FLEX-LasB was performed using an anti-6×His antibody to assess relative expression and recovery (right). (D) Lactate oxidase was expressed in BL21(DE3)pLysS *E. coli* with either N-terminal 6His or FLEX fusion and purified by IMAC.

ExoS is a bifunctional, multidomain *P. aeruginosa* secreted virulence enzyme comprising an N-terminal GTPase-activating protein (GAP) domain and a C-terminal ADP-ribosyltransferase (ADPRT) domain [34, 35]. While truncated constructs of ExoS, particularly the isolated ADPRT domain, have previously been produced recombinantly, expression and purification of full-length ExoS has proven challenging. Residues 51-72 of ExoS have been implicated in membrane association and aggregation, contributing to poor solubility and complex purification behaviour [36]. While the expression of His-FLEX-ExoS yielded detectable soluble protein, purification by IMAC resulted in substantial co-purification of contaminating proteins (Figure 5B, left). Subsequent SEC improved separation of His-FLEX-ExoS but did not resolve the extensive background of co-eluting species (Figure 5B, middle). In contrast, purification by HAC enabled isolation of His-FLEX-ExoS to apparent homogeneity in a single step (Figure 5B, right). Following stringent washes with 1 M NaCl, His-FLEX-ExoS eluted at high ionic strength (2 M NaCl), yielding a single prominent band on SDS-PAGE.

Following this, expression of the secreted *P. aeruginosa* protease LasB, which is typically associated with toxicity and poor expression in *E. coli* [37] was trialled. Recombinant LasB expression in *E. coli* has been widely reported to become incorporated into inclusion bodies [38] or yield only trace amounts of protein that can only be detected by immunoblotting [39]. Consequently, LasB is most commonly purified from culture supernatants of *P. aeruginosa* [40], an approach that is labour-intensive, low-yielding, and inherently variable.

Consistent with these reports, His-tagged LasB was purified at very low levels and could not be purified to homogeneity using IMAC (Figure 5C, left). By contrast, His-LasB-FLEX was isolated at greater concentration and to higher purity in a single HAC step, yielding approximately 0.5 mg of protein per litre of culture (Figure 5C, middle). Western blotting employing an anti 6his allowed us to detect that there was indeed a small quantity of His-LasB post IMAC, but this was substantially lower than the quantity observed for His-LasB-FLEX.

We next examined whether fusion to FLEX altered the enzymatic activity of purified *P. aeruginosa* virulence factors (Supplementary Figure 4). FLEX-tagged ExoU, ExoS, and LasB were assessed using established activity assays for ExoU [22] and ExoS [24] and a newly developed fluorogenic substrate assay for LasB. ExoU activity was measured using a phospholipase A₂ assay, consistent with its function as a patatin-like phospholipase. ExoS activity was quantified using an ADP-ribosyltransferase (ADPRT) assay. For LasB, proteolytic activity was determined using a fluorogenic peptide substrate (Mca-KPLGL-Dpa-AR-NH₂), in which substrate cleavage results in increased fluorescence signal. Following TEV protease cleavage and reverse purification to remove the FLEX tag, kinetic parameters for each enzyme were indistinguishable from those of the uncleaved proteins (Supplementary Figure 4). Together, these data indicate that FLEX fusion does not impair catalytic function and is compatible with proper folding and active-site integrity across mechanistically diverse

1. *P. aeruginosa* virulence enzymes.

As a further case study, the Centre for Automated Protein Research and Innovation (CAPRI), a protein production facility at the University of Liverpool, evaluated FLEX in a challenging industrial expression project involving L-lactate oxidase from *Aerococcus viridans*.

Following IMAC purification, conventionally His-tagged lactate oxidase was recovered at low yield and co-eluted with substantial impurities (Figure 4D). In contrast, incorporation of the FLEX tag-applied after IMAC resulted in markedly improved recovery and purity of the target protein. These findings provide an independent validation of FLEX performance in a real-world protein production setting and highlight its potential utility for difficult-to-express enzymes in both academic and contract manufacturing workflows.

### Compatibility FLEX with mammalian expression systems

Mammalian expression systems typically exploit peptide-based tags, including epitope tags (e.g. Myc, HA, and FLAG) and Strep tag, that are captured from lysates using immobilised antibodies or, in the case of Strep tag, streptavidin derivatives. Recombinant proteins are typically eluted using excess peptide (or Desthiobiotin or biotin for Strep-based tags) or alternatively through engineering of site-specific proteolytic cleavage sites to release of the POI from the immobilised resin. Antibody-based affinity chromatography can also utilise a low pH elution, but this may have adverse effects on the fusion partner. A major limitation for these approaches that has impeded their widespread adoption in high throughput protein purification workflows is the relatively high costs of the affinity/immunoprecipitation resins. This expense is exacerbated by the fact that these are typically ‘single use’ resins or are challenging to efficiently regenerate for multiple rounds of purification. In contrast, the chemically robust heparin-based affinity resins can be regenerated by washing in high-salt concentrations. Co-purification of contaminant proteins is also an important consideration that can confound the analytical interpretation of mammalian cell-derived protein function and must be carefully controlled. This particularly makes polyHis tags unsuitable for mammalian cell protein purification due to the propensity of chelated metal matrices to non-specifically bind proteins resulting in high background noise. Owing to their large size, MBP and GST tags, which are primarily optimised for bacterial expression, can interfere with the structure, subcellular location and the biological functions of their fusion partner and may therefore block critical post-translational modification [41–43]. These factors have driven a preference for smaller affinity tags for mammalian cell applications.

We next tested whether FLEX could function as a purification handle in mammalian cells. His-FLEX-tagged human protein kinase A (PKA) kinase domain was transiently expressed in HEK293T cells and purified from cell lysates by either nickel or heparin affinity chromatography. Reassuringly, we detected robust expression of PKA in HEK293T cell lysates (Figure 6A). While IMAC resulted in significant co-purification of host proteins, a single heparin affinity step yielded a strikingly pure preparation of PKA (Figure 6A), demonstrating that FLEX is compatible with mammalian expression and enables high-purity purification without the need for multistep workflows.

**Figure 6.**
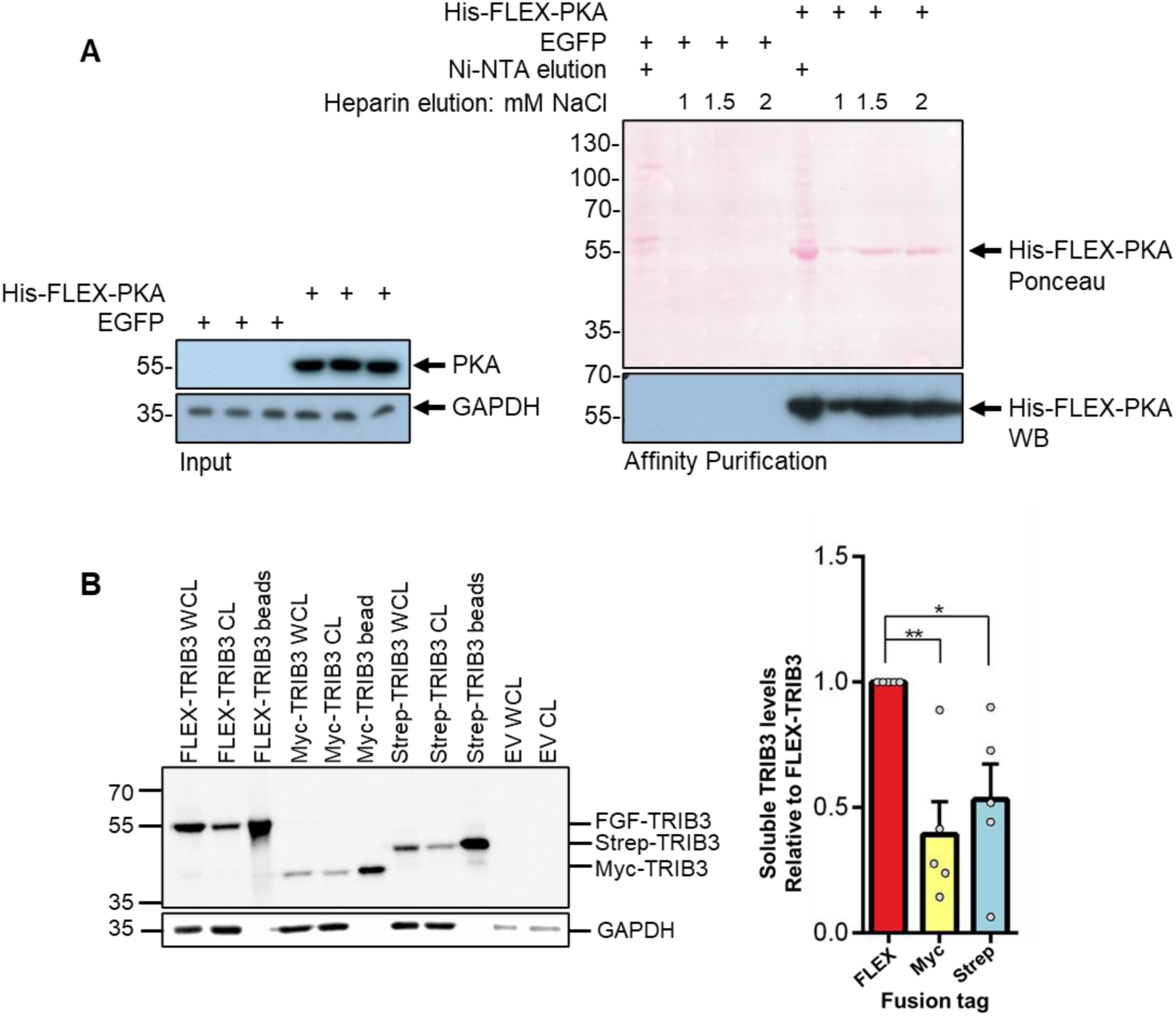
FLEX tag improves recovery and is compatible with mammalian expression systems. (A) The kinase domain of protein kinase A (PKA) was expressed in HEK293T cells as an N-terminal His-FLEX fusion from a pTWIST CMV mammalian expression vector. Clarified lysates were subjected to pulldown using either Ni-NTA (IMAC) or heparin affinity resin to compare purification efficiency of the His–FLEX construct by metal versus heparin binding. (B) TRIB3 was expressed as N-terminal FLEX-, Myc-, or Strep-tagged constructs in HEK293T cells to compare FLEX with commonly used affinity tags. Clarified lysates were incubated with heparin, anti-Myc, or Strep-avidin resins, respectively, followed by washing in 500 mM NaCl Tris buffer and elution heating (98°C) of resins in an equal volume (10 ml) of 1x Laemmli sample buffer for 10 minutes, before centrifugation to collect bound TRIB3. Densitometry was quantified for clarified lysates of FLEX-TRIB3, Myc-TRIB3, and Strep-TRIB3 using ImageJ and plotted using GraphPad Prism 6, the student’s T test was used to determine statistically significant differences from five independent experiments. For all experiments, HEK293T cells were transiently transfected with plasmid DNA using polyethylenimine and harvested 40-48 h post-transfection. Cells were lysed in Tris-based buffer containing NaCl, Triton X-100, glycerol, DTT, and protease/phosphatase inhibitors, followed by sonication and clarification by centrifugation. Samples were analysed by SDS-PAGE and immunoblotting using anti-His, anti-TRIB3, or control antibodies.These data show that FLEX tagging improves recovery of challenging mammalian proteins and enables efficient purification compared with conventional affinity tags. WCL, whole-cell lysate; CL, clarified lysate.

#### FLEX improves TRIB3 expression compared to gold standard mammalian expression tags

TRIB3 is a stress-responsive pseudokinase that regulates multiple signalling pathways involved in the control of cell survival in diverse cell types, with key roles in cancer and metabolic disorders [44]. It is an extremely challenging protein to express and current approaches allow only limited purification of TRIB3, necessitating further optimization to enable robust biochemical analyses and scalable inhibitor screening analogous to those established for TRIB1 and TRIB2 [45–47]. The FLEX affinity tag was compared with the widely used Strep and Myc tags using small-scale pulldown assays following transient transfection in HEK-293T cells. Immunoblot analysis showed consistently higher protein levels of total and soluble TRIB3 when tagged with FLEX. The recovered FLEX-TRIB3 bound to heparin beads was also greater than that recovered from the more commercially expensive, anti-Myc resin (for Myc-TRIB3) and strep-avidin resin (for Strep-TRIB3).

Together, the TRIB3 and PKA pulldown experiments demonstrate that FLEX can enhance both expression and recovery of difficult proteins from mammalian expression systems, providing a practical route to obtaining sufficient material for downstream biochemical characterisation and inhibitor screening.

## Discussion

The expression and purification of unstable, aggregation-prone, or cytotoxic recombinant proteins remains a persistent challenge across structural biology, enzymology, and translational research. Fusion tags are widely used to improve solubility and enable affinity purification, yet most existing tags require trade-offs between expression yield, purification stringency, tag size, and downstream compatibility. In this study, we describe the rational engineering and validation of FLEX, a compact heparin-binding fusion tag derived from human FGF1 that combines enhanced stability with high-affinity, high-stringency purification capability.

### Structure-guided stabilisation of an FGF1-derived scaffold

The intrinsic instability of wild-type FGF1 has historically precluded its use as an affinity ligand or fusion partner, despite its strong heparin-binding capacity. By combining targeted truncations with structure-guided mutations, we were able to dramatically improve both thermal and chemical stability. FLEX exhibited an increase in melting temperature of more than 20 °C and resistance to denaturation even at 6 M urea (Figure 4), establishing it among the more stable fusion tags reported to date. Improved stability was accompanied by enhanced expression yield, which in turn increased the fraction of correctly folded, affinity-competent material available for purification.

Importantly, stabilisation was achieved without compromising glycosaminoglycan recognition or the characteristic β-trefoil fold. Heparin recognition by FGF1 is mediated by a spatially distributed electropositive surface formed by multiple Lys and Arg residues across the folded β-trefoil core, rather than a single linear binding motif (Figure 1). The enhanced affinity observed for FLEX therefore appears to arise from expansion and optimisation of the positively charged surface surrounding the binding interface, rather than direct modification of the canonical interaction site. This distinction is critical, as it preserves the predictable binding mode of FGF1-heparin interactions while improving performance. Together, these findings indicate that heparin binding by FGF1 is governed by surface electrostatics and conformational stability, both of which are optimised in the engineered FLEX scaffold.

### FLEX in the context of existing fusion tags

A wide range of fusion tags have been developed to address protein solubility and purification challenges, yet no universal solution exists. Small affinity tags such as polyhistidine (His-tags) are minimally invasive but often suffer from non-specific binding and limited purification stringency, particularly for proteins expressed at low levels or in complex lysates such as those derived from mammalian expression systems. Larger solubility-enhancing tags, including maltose-binding protein (MBP), glutathione S-transferase (GST), NusA, and thioredoxin, can improve expression but frequently introduce new challenges, including proteolytic instability, inefficient cleavage, interference with folding or function, and complications for structural studies due to their large size.

FLEX occupies an intermediate and underexplored design space: a moderately sized (∼15.5 kDa), intrinsically stable, globular fusion tag that enables high-affinity, non-metal-based purification. Unlike MBP or GST, FLEX does not rely on carbohydrate binding or enzymatic activity, and unlike His-tags, it supports purification under highly stringent conditions.

Immobilised heparin matrices could be washed with 1000 mM NaCl without loss of bound FLEX-tagged protein, enabling efficient removal of host contaminants that commonly co-purify during IMAC.

Heparin affinity chromatography is widely used for purification of growth factors, nucleic acid-binding proteins, and viral components, but has rarely been exploited as a general-purpose fusion tag strategy. One limitation has been the relatively modest affinity of many heparin-binding proteins, which restricts washing conditions and leads to co-purification of host proteins with similar charge properties. The enhanced affinity of FLEX for immobilised heparin, quantified by ITC and reflected in delayed elution at high ionic strength (Figure 3), overcomes this limitation. FLEX-tagged proteins eluted only at 1.5-2.0 M NaCl (Supplementary Figure 2), providing a level of selectivity that rivals or exceeds IMAC, amylose-, or glutathione-based systems, while avoiding competitive eluents.

This high-stringency purification capability simplifies workflows and avoids potential complications associated with metal chelation, imidazole removal, or interference with downstream biochemical and biophysical assays. Consistent with this, FLEX demonstrated broad assay compatibility with minimal impact on protein activity (Supplementary Figure 4), supporting its use as a general-purpose fusion partner for challenging recombinant proteins across expression platforms.

### FLEX enables expression of challenging proteins

A major strength of FLEX is its performance across a diverse set of challenging proteins, including bacterial toxins and secreted proteases. In several cases, most notably ExoU, LasB, and full-length ExoS, conventional fusion tags failed entirely, yielding either insoluble protein, extensive degradation, or recovery of tag alone. The successful expression and purification of these proteins as FLEX fusions suggests that the stabilising properties of the tag extend beyond simple affinity handling. The asymmetric surface properties of FLEX, with a hydrophilic heparin-binding face and a more hydrophobic opposing surface, raise the possibility that FLEX may reduce aggregation by shielding exposed hydrophobic regions of fusion partners during folding and expression. While this hypothesis was not directly tested here, it is consistent with the observed improvements in soluble yield and tractability across multiple unrelated targets.

### Compatibility with mammalian expression systems

An additional advantage of FLEX is its compatibility with mammalian expression platforms. Heparin affinity purification is routinely used for secreted and cell-surface proteins in mammalian systems yet has rarely been integrated into general fusion tag strategies. The ability to purify His-FLEX-tagged PKA to near homogeneity from HEK293T lysates in a single step demonstrates that FLEX can function effectively in complex eukaryotic proteomes, where IMAC often suffers from high background binding due to histidine-rich host proteins and metal-binding contaminants. The high-stringency washing enabled by FLEX-heparin interactions allows efficient removal of mammalian host proteins while preserving yield, simplifying workflows that would otherwise require multiple chromatographic steps.

Strikingly, FLEX also improved recovery of the stress-responsive pseudokinase TRIB3, a protein that has historically proven difficult to express and purify despite extensive optimisation efforts in the TRIB field. Increased levels of soluble TRIB3 in HEK293T cells suggest that FLEX can enhance folding or stability in the eukaryotic intracellular environment, potentially by reducing exposed hydrophobic patches, shielding aggregation-prone surfaces, or stabilising partially folded intermediates during translation. This mirrors the stabilising effects observed in bacterial expression and indicates that FLEX may act not only as an affinity handle but also as a chaperone-like scaffold across expression hosts.

The compatibility of FLEX with mammalian systems has several practical implications. Many recombinant protein workflows, such as those used in kinase inhibitor screening, antibody development, and virulence factor studies relevant to *Pseudomonas aeruginosa* toxins like ExoU, require initial production in *E. coli* followed by expression of validated constructs in mammalian cells for functional or structural studies. A single tag that performs consistently across these platforms reduces construct redesign, accelerates optimisation cycles, and facilitates translation from biochemical discovery to cellular validation. In addition, heparin-based purification avoids metal chelation steps that can interfere with downstream assays, particularly those sensitive to divalent cations or redox-active metals. Although not all mammalian proteins will benefit equally, particularly those lacking accessible FLEX surfaces or those sensitive to high ionic strength elution, the results presented here highlight the unexpected breadth of FLEX utility in eukaryotic systems.

Future work will explore its performance with secreted proteins, membrane-associated targets, and stable cell-line production, as well as its compatibility with automated purification pipelines.

### Limitations and future directions

While FLEX provides several practical advantages, certain limitations should be acknowledged. Purification via heparin affinity chromatography typically requires elution at high ionic strength, which may necessitate buffer exchange prior to downstream applications and could be incompatible with proteins that are particularly sensitive to elevated electrolyte concentrations. To mitigate this, FLEX constructs were designed with an intervening TEV protease site to enable tag removal and reverse purification under milder conditions, as demonstrated for ExoU and for short inhibitor peptides. In cases where high-salt exposure remains problematic, optimisation of wash and elution conditions or integration of orthogonal purification steps may be required.

Although FLEX performed robustly across a diverse panel of challenging proteins, it is unlikely to be universally optimal. Systematic benchmarking against established solubility and affinity tags across broader protein classes and expression hosts will be important to define its performance envelope. We also observed context-dependent differences between heparin affinity chromatography and immobilised metal affinity chromatography for His-FLEX constructs. While heparin resin generally enabled high-purity, single-step purification, in some cases higher recovery was achieved by IMAC. Such behaviour likely reflects differences in resin capacity, ligand density, and accessibility of the affinity motifs within individual fusion proteins. Steric occlusion of the heparin-binding surface altered electrostatic presentation, or partial engagement of FLEX in stabilising intramolecular interactions with the fusion partner could all reduce effective binding to immobilised heparin. Similar context dependence has been reported for other fusion partners, where tag performance varies with domain architecture, linker design, and oligomeric state.

Future work will focus on defining the mechanistic basis of FLEX-mediated stabilisation, including its effects on co-translational folding, aggregation suppression, and intracellular trafficking in both bacterial and mammalian systems. Extending FLEX to periplasmic, secretory, and membrane-protein expression platforms will further test its generality, particularly for disulfide-bonded or extracellular targets. Structural and biophysical characterisation of FLEX-fusion constructs, combined with kinetic folding assays and ribosome-profiling approaches, should clarify how FLEX modulates stability and affinity presentation across different protein contexts. Together, such studies will help refine design rules for deploying FLEX in recombinant protein production pipelines and identify cases where alternative strategies remain preferable.

### Conclusions

In summary, FLEX represents a new class of fusion tag that combines structure-guided stabilisation with high-affinity heparin binding to enable high-purity, high-yield production of difficult recombinant proteins. By addressing limitations inherent to both small affinity tags and large solubility tags, FLEX offers a practical and broadly applicable solution for protein expression and purification in both academic and industrial settings.

## Acknowledgments

This work was supported by a UKRI Impact Accelerator Award (2022-2025), Fight for Sight (UK) research grants 5175 and 5176, and the Biotechnology and Biological Sciences Research Council (BBSRC; BB/X002780/1). We thank the CAPRI Protein Production Service at the University of Liverpool for technical support and assistance with recombinant protein expression.

---

**Supplementary Figure 1:**
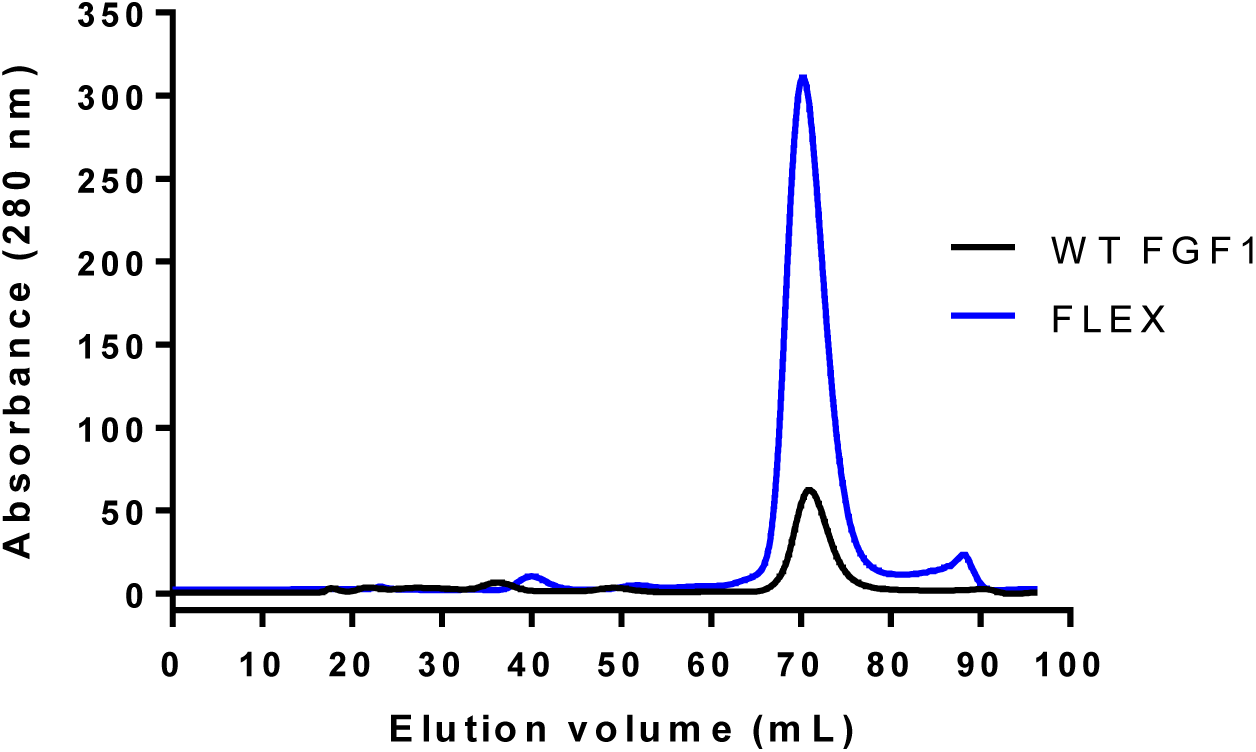
Size-exclusion chromatography (SEC) profiles of His-tagged WT FGF1 (black) and FLEX (blue) following immobilised metal affinity chromatography (IMAC). SEC using a Superdex 200 16/600 column was performed to assess the oligomeric state of the purified proteins prior to downstream analyses. Elution volume is shown with protein detected by absorbance at 280 nm. Both His-WT FGF1 and His-FLEX eluted as monomeric species corresponding to the predicted 20 kDa molecular weight.

**Supplementary Figure 2:**
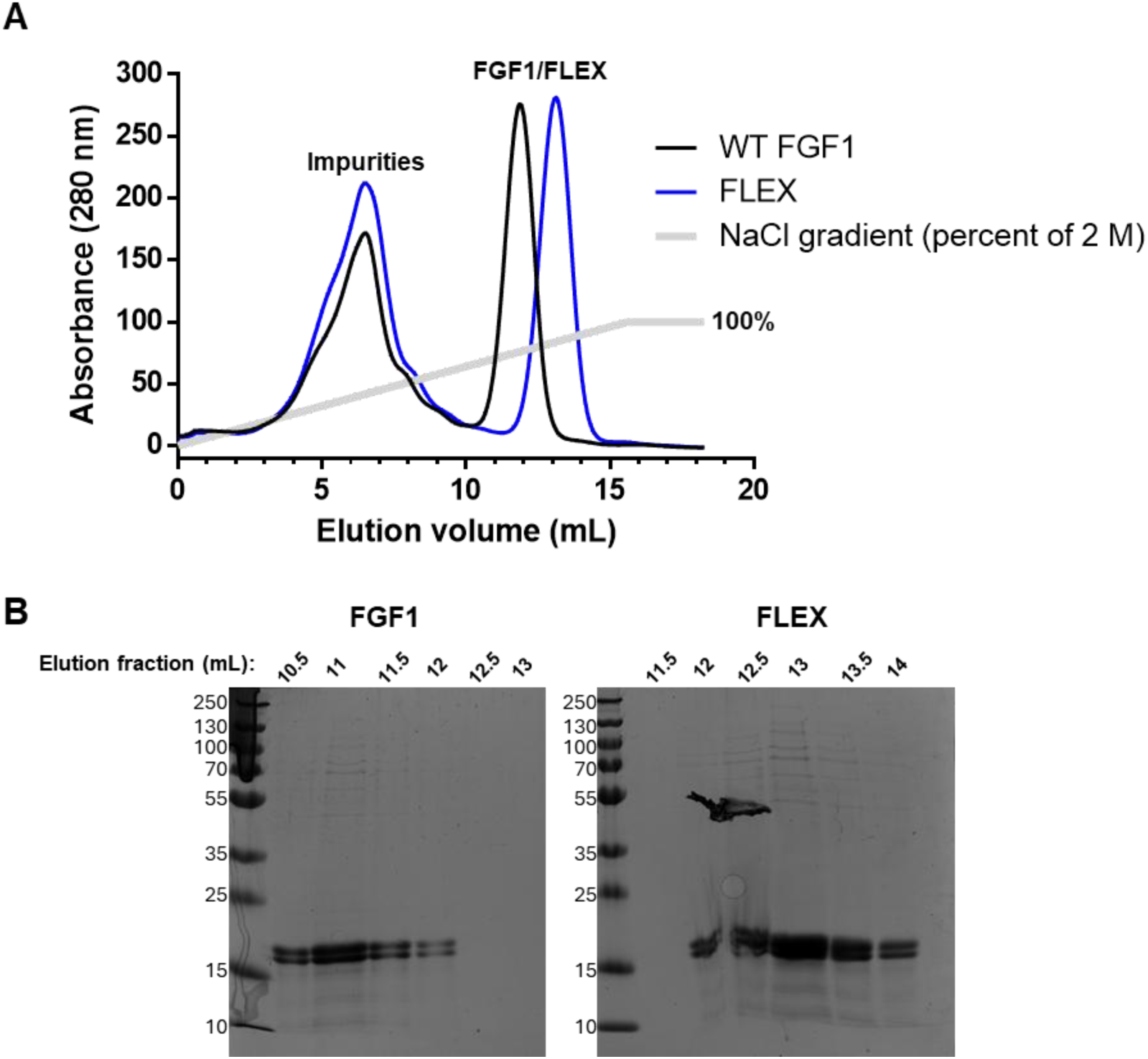
Heparin affinity chromatography of WT FGF1 and FLEX using a linear NaCl gradient. Clarified *E. coli* lysates were applied to immobilised heparin columns, washed with 300 mM NaCl, and eluted with a 300 mM-2 M NaCl gradient. Protein elution was monitored by absorbance at 280 nm. The NaCl gradient is indicated by the gray line. (B) Eluted fractions containing FGF1 and FLEX were analysed by SDS-PAGE.

**Supplementary Figure 3:**
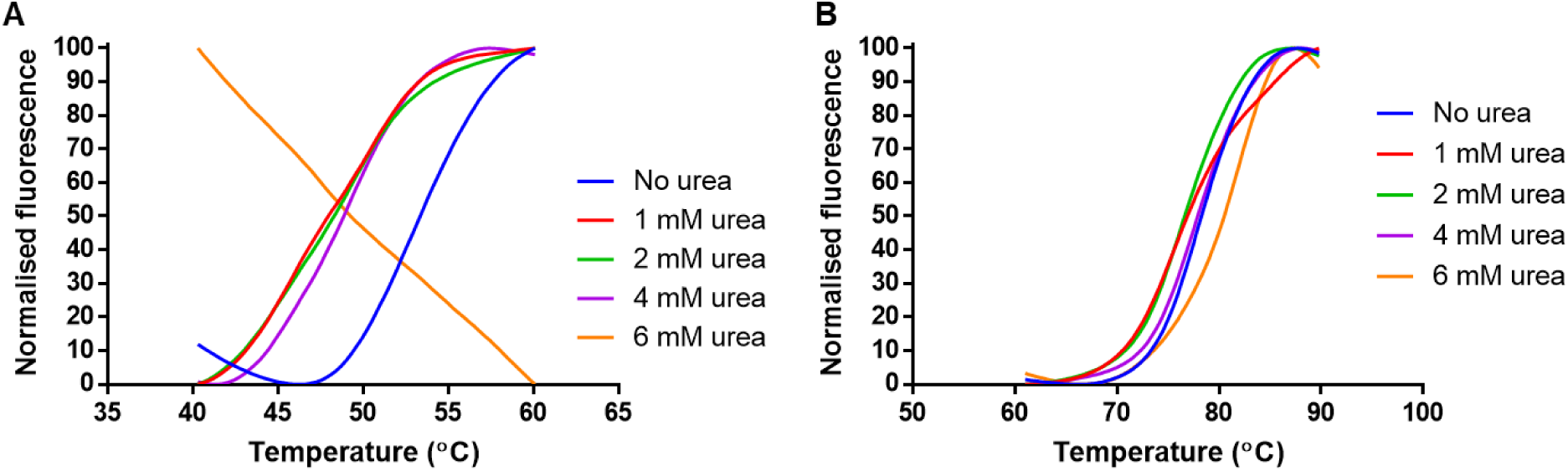
Thermal denaturation profiles of WT FGF1 and FLEX under chemical denaturation. (A) Representative DSF unfolding curves for WT FGF1 in the presence of increasing concentrations of urea (1-6 M). (B) Representative DSF unfolding curves for FLEX under identical urea conditions.

**Supplementary Figure 4:**
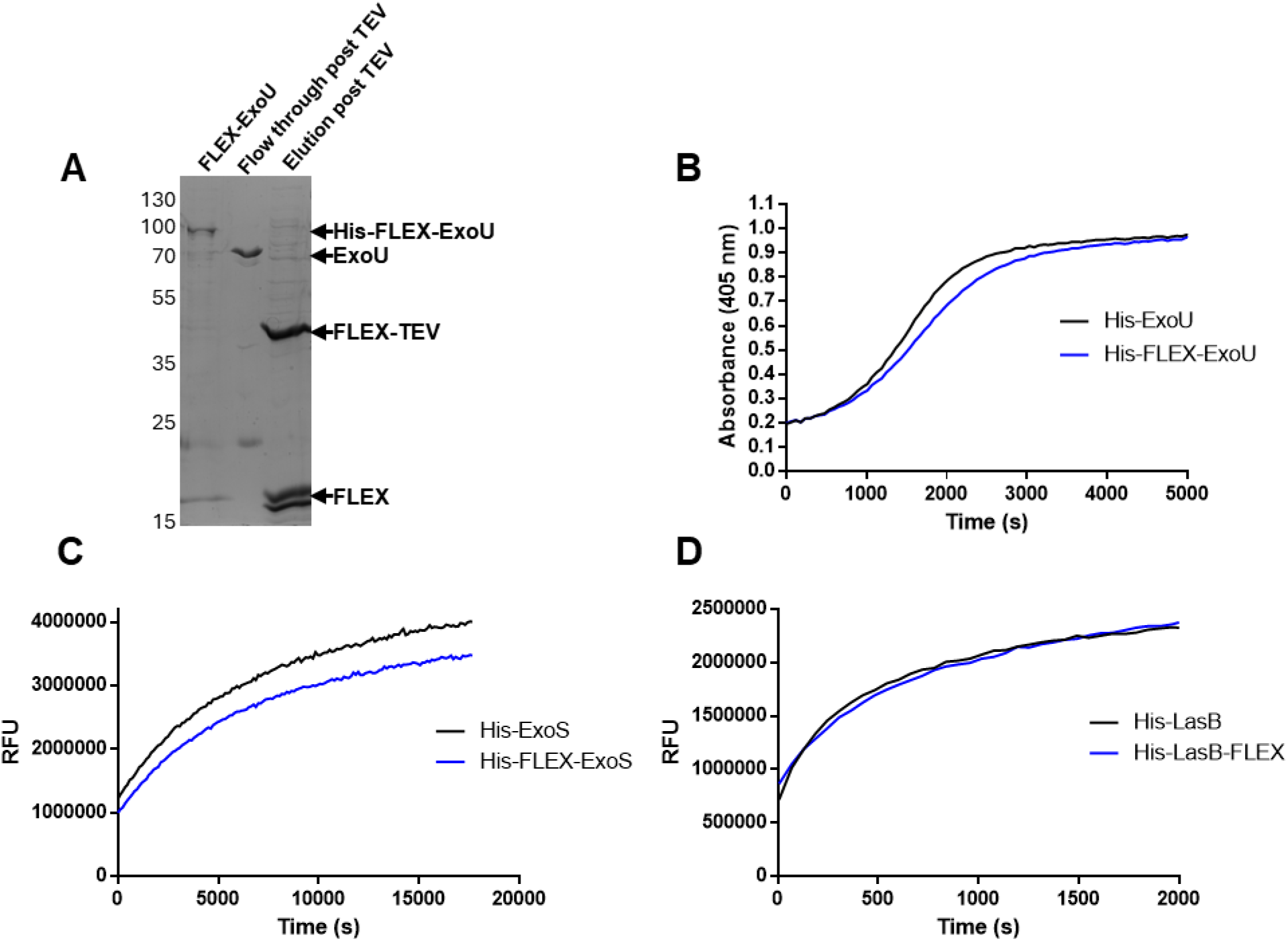
Enzymatic activity of His- and FLEX-tagged *P. aeruginosa* virulence factors. (A) Removal of the FLEX tag. His-FLEX fusion proteins were treated with His-FLEX tagged TEV protease and incubated with heparin resin to remove His-FLEX and His-FLEX-TEV, yielding tag-free enzymes in the flow through by reverse affinity purification. An elution of 2M NaCl was employed to remove the His-FLEX tag and His-FLEX-TEV from the heparin resin. Activities of purified ExoU (B), ExoS (C), and LasB (D) were assessed using established biochemical assays. Enzymes were used at 100 nM. Where indicated, the FLEX tag was removed by TEV protease cleavage followed by reverse affinity purification prior to analysis. (A) Phospholipase activity of ExoU measured using arachidonyl thio-phosphatidylcholine as substrate, in the presence of 1 µM phosphatidylinositol-4,5-bisphosphate (PIP₂) and 5 µM mono-ubiquitin for activation. Hydrolysis was monitored using 5,5′-dithiobis-(2-nitrobenzoic acid) (DTNB) as a reporter, with an increase in absorbance at 405 nm. Assays were performed in Tris buffer (pH 8.2) containing 100 mM NaCl. (B) ADP-ribosyltransferase activity of ExoS assessed by auto-ADP-ribosylation using ε-NAD (25 µM) as substrate and 5 µM 14-3-3β as an activating co-factor, in Tris buffer (pH 8.2) containing 100 mM NaCl. (C) Proteolytic activity of LasB measured using the fluorogenic peptide substrate Mca-KPLGL-Dpa-AR-NH₂ (25 µM) in Tris buffer (pH 8.2) containing 100 mM NaCl, 10 µM ZnSO₄, and 100 µM CaCl₂.

